# Coordination of Glucose and Glutamine Metabolism in Tendon is Lost in Aging

**DOI:** 10.1101/2024.12.19.629426

**Authors:** Samuel J. Mlawer, Felicia R. Pinto, Katie J. Sikes, Brianne K. Connizzo

**Author notes:** Correspondence: Brianne K. Connizzo 44 Cummington Mall Boston, MA, 02215, United States E. Co-first author (Samuel J. Mlawer, Felicia R. Pinto).

## Abstract

Tendinopathy is an age-associated degenerative disease characterized by a loss in extracellular matrix (ECM). Since glucose and glutamine metabolism is critical to amino acid synthesis and known to be altered in aging, we sought to investigate if age-related changes in metabolism are linked to changes in ECM remodeling. We exposed young and aged tendon explants to various concentrations of glucose and glutamine to observe changes in metabolic processing (enzyme levels, gene expression, etc.) and matrix biosynthesis. Interestingly, we found that glutamine processing is affected by glucose levels, but this effect was lost with aging. ECM synthesis was altered in a protein-dependent manner by increased glucose and glutamine levels in young tendons. However, these changes were not conserved in aged tendons. Overall, our work suggests that glucose and glutamine metabolism is important for ECM homeostasis, and age-related changes in nutrient metabolism could be a key driver of tendon degeneration.

## Introduction

Thirty-three million musculoskeletal injuries are reported annually, with roughly 50% consisting of injuries to soft tissue including tendon.^1^ Incidences of tendon injury and degeneration are even more prevalent in the aged population, with rotator cuff tendinopathy affecting over 50% of the population above the age of 60. Our previous work has identified that aged tendons have deficiencies in extracellular matrix (ECM) maintenance, but the exact cellular mechanisms that contribute to this dysfunction are still not well understood.^2–4^ There are many hallmarks of cell aging that could influence ECM synthesis and breakdown and thus contribute to the development of tendinopathy.^5^ One of these factors of aging is mitochondrial dysfunction, where the ability to produce ATP from glucose through cellular respiration is impaired.^5,6^ Our previous work has found decreased metabolic activity through cellular respiration in aged tendons compared to young tendons.^3,4^ However, it remains to be seen if this altered metabolism in aged tendons directly leads to our observed ECM remodeling deficiencies. Glucose and glutamine are two key metabolites that are involved in energy production, as well as in the synthesis of key amino acids for ECM proteins, and therefore could be a potential link between age-related metabolic and ECM changes.^7–10^

While glucose is the main source of energy for all cells, few studies have observed how tenocytes directly utilize glucose, as they are highly quiescent at homeostasis.^11^ However, clinical studies have found that patients with type II diabetes mellitus and corresponding elevated blood glucose levels, are four times as likely to develop tendinopathy.^12^ However, little is known about how these atypical levels of glucose in healthy tendon influence tendon homeostasis and the progression of tendon disease. Non-diabetic *in vivo* models have observed a positive response with a high glucose diet leading to increased tendon thickness and stiffness following injury.^13^

Previous work in tendon cells has observed the opposite, with high glucose repressing cell viability, increasing apoptosis and inflammation, and decreasing expression of gene markers for matrix synthesis and fibrillogenesis.^14–17^ Clinical studies typically have an older patient population whereas laboratory studies focus on young or clonal cell populations, and thus age may be a confounding variable in drawing conclusions regarding the role of glucose in tendon health. It’s also important to consider other nutrients that might be relevant for ECM synthesis, such as glutamine.

Glutamine is the most abundant amino acid in the body and is vital for a variety of cell processes, including amino acid synthesis, protein synthesis, nucleotide synthesis, and redox homeostasis.^18^ Importantly for tendon, glutamine is synthesized into proline, which together with its derivative, hydroxyproline, make up approximately 23% of amino acids in collagen.^18,19^ Tendon structure is primarily composed of collagen, making up 60-85% of tendon dry weight.^20^ Since collagen synthesis is promoted by proline, an understanding of the effects of glutamine on tendon homeostasis is important. Despite glutamine being a common addition to media for tendon studies, its effects in tendon explants have not been studied. Previous studies in fibroblasts have found that glutamine supplementation in media upregulates collagen synthesis and non-collagenous protein synthesis.^18,21,22^ On the other hand, other studies that looked at higher concentrations of glutamine found that excess glutamine instead suppressed collagen synthesis.^23^ This means that a proper level of glutamine is vital to ECM homeostasis, since both low and high concentrations can influence synthesis. Glutamine can also be transported to the mitochondria and used to create α-ketoglutarate, which is then used in the citric acid cycle for energy production in coordination with glucose. Since there is a relationship between these pathways and previous studies have observed crosstalk between the effects of glucose and glutamine in cells, it is important to understand not only their individual effects, but also the coordination between their effects in tendon.^24–27^

The overall goal of our studies is to delineate the contribution of metabolic changes to age-related dysfunction in ECM homeostasis. Since *in vivo* tools to quantify protein-specific turnover in real-time are not yet available and isolated culture removes cells from their native microenvironment, it has been historically quite challenging to link changes in cell biology with ECM remodeling. To address, this we capitalize on our established *ex vivo* explant model that enables the study of real-time ECM synthesis and breakdown, as well as simultaneous interrogation of live cells maintained in a physiologically-relevant three-dimensional environment.^4^ Furthermore, these tissues can be harvested from young and old animals allowing for study of aging processes directly. The purpose of the present study was to investigate the response of young and aged tendon explants to media supplementation with various concentrations of glucose and glutamine. We hypothesized that elevated levels of glucose and glutamine would lead to increased ECM remodeling in young explants. Based on previous studies, we also hypothesized that aged explants would have a muted response to the elevated glucose and glutamine levels.

## Results

### Coordination of glucose and glutamine processing is lost in aged tendons

We first looked at how tendon explants’ uptake of glucose and glutamine varied with medium glucose and glutamine concentration. Tendon glucose uptake increased with high glucose medium compared to low glucose medium in both age groups. In addition to the expected glucose-dependent response, there was also a glutamine-dependent response; increased glucose uptake in our high glucose groups was only present when there was also glutamine supplementation. The highest glucose uptake occurred with 200 µM glutamine (Figure 1A). Glutamine uptake was also dependent on both glucose and glutamine levels. In both age groups, glutamine uptake was greatly stimulated with the combination of high glucose and high glutamine. Surprisingly, in aged explants we found that the combination of low glucose and high glutamine led to negative uptake values, which suggests that there was more glutamine in the medium than originally added (Figure 1B). This suggests decreased glutamine uptake of course, but also potentially that tenocytes secreted additional glutamine under these conditions.

**Figure 1.**
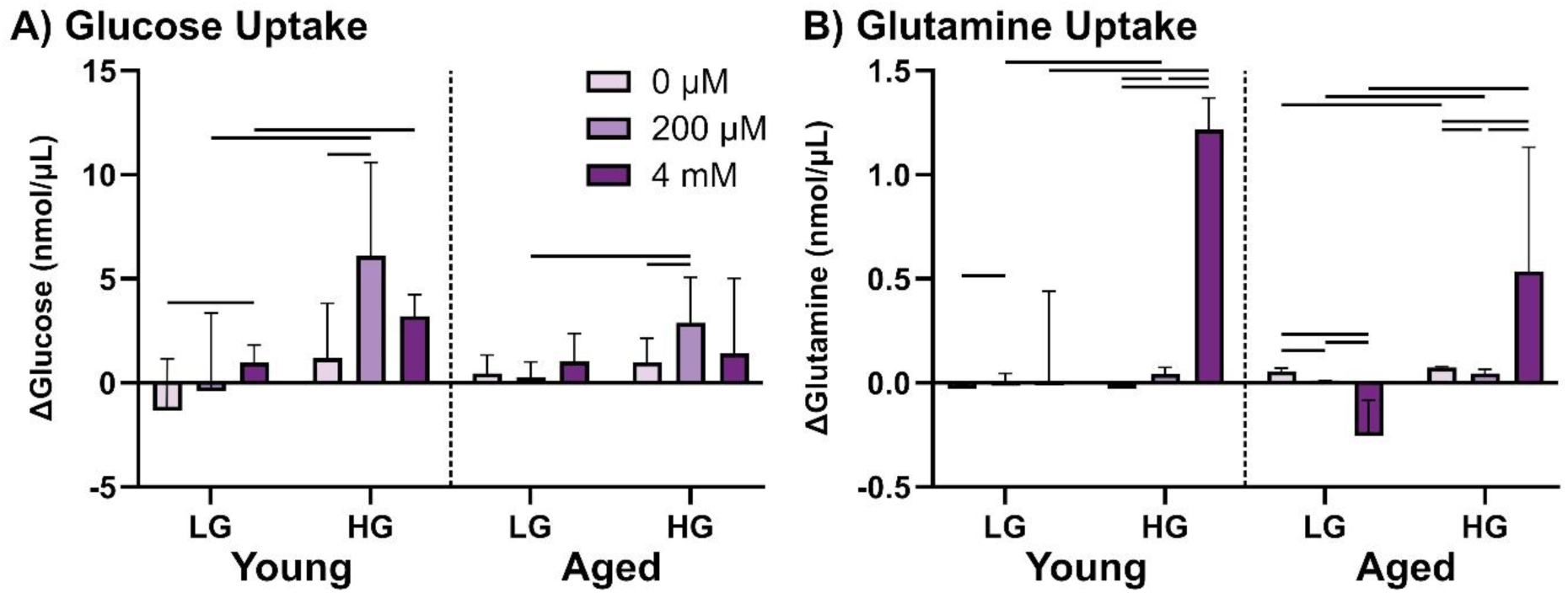
(A) Glucose and (B) Glutamine uptake by young and aged explants at Day 7 as a function of both glucose and glutamine media supplementation. Data is presented as mean ± 95% confidence interval. Significance between media groups is denoted with a bar (-) with p<0.05. [LG = low glucose, HG = high glucose, 0 µM = 0 µM glutamine, 200 µM = 200 µM glutamine, 4 mM = 4 mM glutamine]

We next were interested in how these changes in metabolite concentration affected gene expression of certain genes on the glucose and glutamine pathways. Gene expression of pyruvate dehydrogenase kinase 2 (PDK2) and glutathione synthetase (GSS) was increased by high glucose in aged explants only (Figure 2A). We also found that high glucose in young explants increased expression of the glutamine pathway genes glutaminase (GLS), glutamate dehydrogenase 1 (GLUD1), solute carrier family 1 member 5 (SLC1A5), and glutamic-oxaloacetic transaminase (GOT1). High glutamine concentrations increased expression of the glucose pathway genes PDK2 and phosphoenolpyruvate carboxykinase 1 (PCK1) in aged tendons only. Among glutamine-related genes, we found increased expression of GLUD1 with our highest glutamine concentration in aged tendons only. In young explants, SLC1A5 expression was decreased in 200 μM and 4 mM glutamine groups (Figure 2B). We then looked at how these changes in gene expression affected protein level of a few key enzymes involved in the glutamine pathways, which were chosen based on gene expression data. We found that glutathione (GSH) decreased with high glucose in young, with no such change in aged explants (Figure 3A). The activity of glutamate dehydrogenase (GDH) decreased with high glucose in aged explants, with no change in young (Figure 3B).

**Figure 2.**
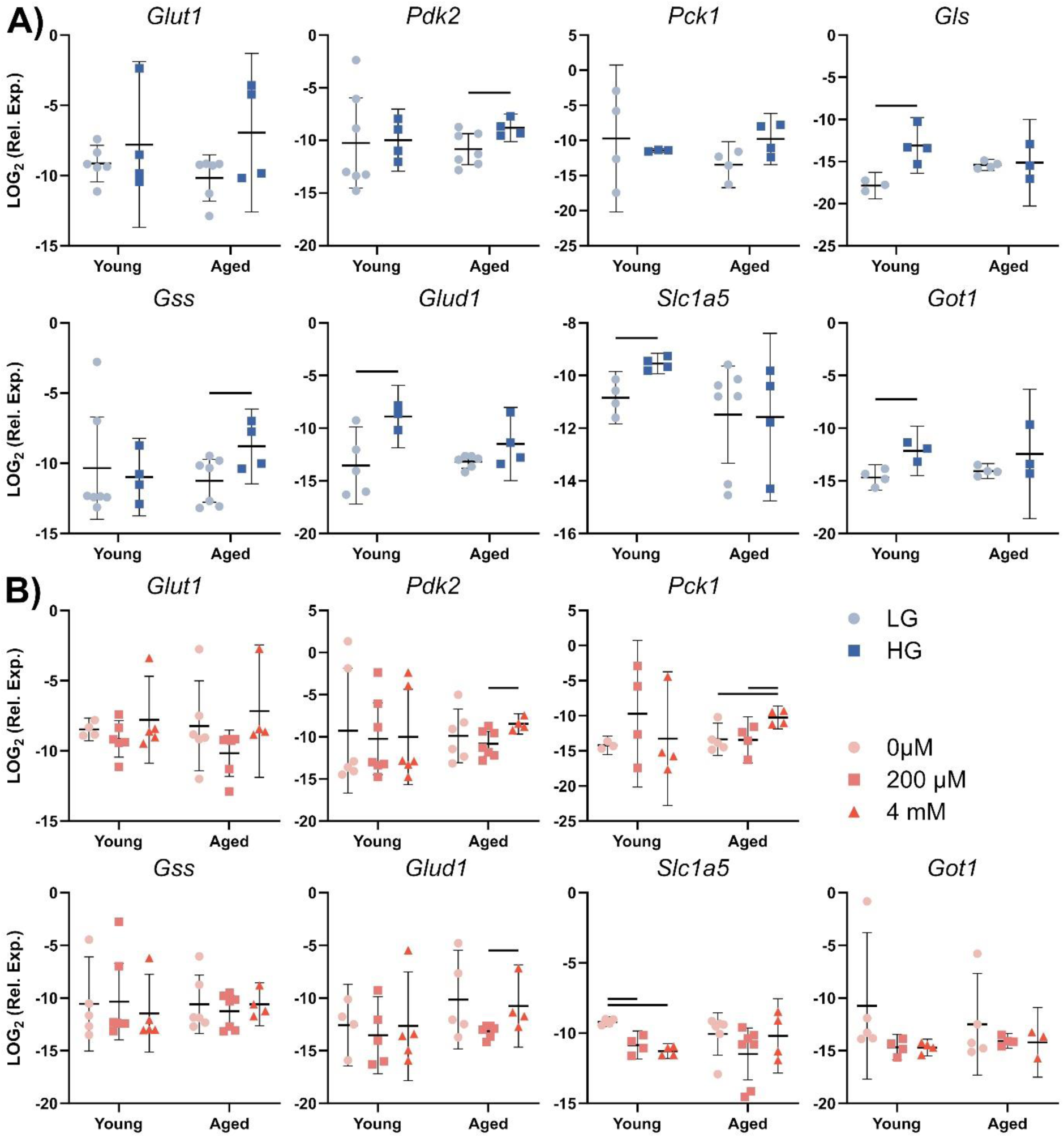
Measurement of mRNA levels of glucose and glutamine pathway genes for young and aged explants as a function of (A) glucose and (B) glutamine concentration. Data is presented as mean ± 95% confidence interval. Significance between media conditions is denoted with a bar (-) with p<0.05. [LG = low glucose, HG = high glucose, 0 µM = 0 µM glutamine, 200 µM = 200 µM glutamine, 4 mM = 4 mM glutamine]

**Figure 3.**
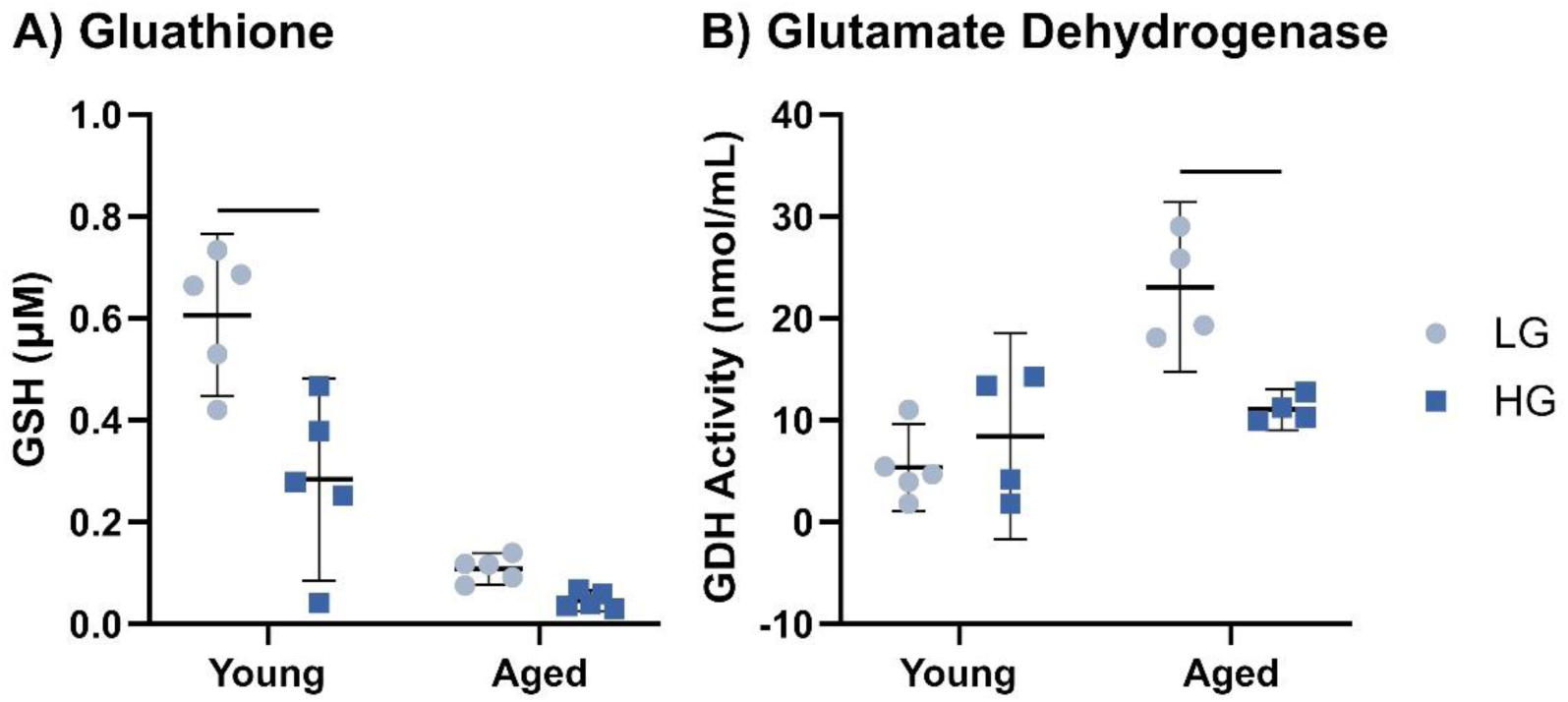
Measurement of protein levels of (A) glutathione (GSH) and (B) glutamate dehydrogenase (GDH) for young and aged explants as a function of glucose concentration with glutamine concentration at 200 μM. Data is presented as mean ± 95% confidence interval. Significance between media conditions is denoted with a bar (-) with p<0.05. [LG = low glucose, HG = high glucose]

### Glucose- and glutamine-dependent effects on ECM synthesis are lost in aged tendons

We then sought to investigate how medium glucose and glutamine concentration influenced tenocyte and ECM homeostasis. Metabolism and DNA content significantly increased with glucose in the young group, while there was no change in either in aged tendons (Supplemental Figure S1A & S1B). Tendon sGAG content and synthesis decreased in young samples with increased glucose, with no differences in aged tendons (Fig. 4A & 4B). We also observed increased expression of both decorin and biglycan with high glucose (Figure 4C & 4D). We observed different effects with glutamine supplementation, with no change in either metabolism or DNA content (Supplemental Figure S1A & S1B). While sGAG content saw no significant changes with glutamine, sGAG synthesis was glutamine dependent. In young groups, 200 µM of glutamine stimulated sGAG synthesis, while 4 mM made no difference compared to tendons with no added glutamine. However, aged tendons did not experience the same response to our highest glutamine concentration, and instead sGAG synthesis increased with increased glutamine (Figure 4A & 4B). In addition, biglycan expression was sensitive to glutamine concentration in young explants, with reduced expression in the no glutamine group (Figure 4D). Interestingly, these changes to proteoglycan gene expression were not accompanied by any changes to matrix metallopeptidase 3 (MMP-3) gene expression in either age group (Figure 4E).

**Figure 4.**
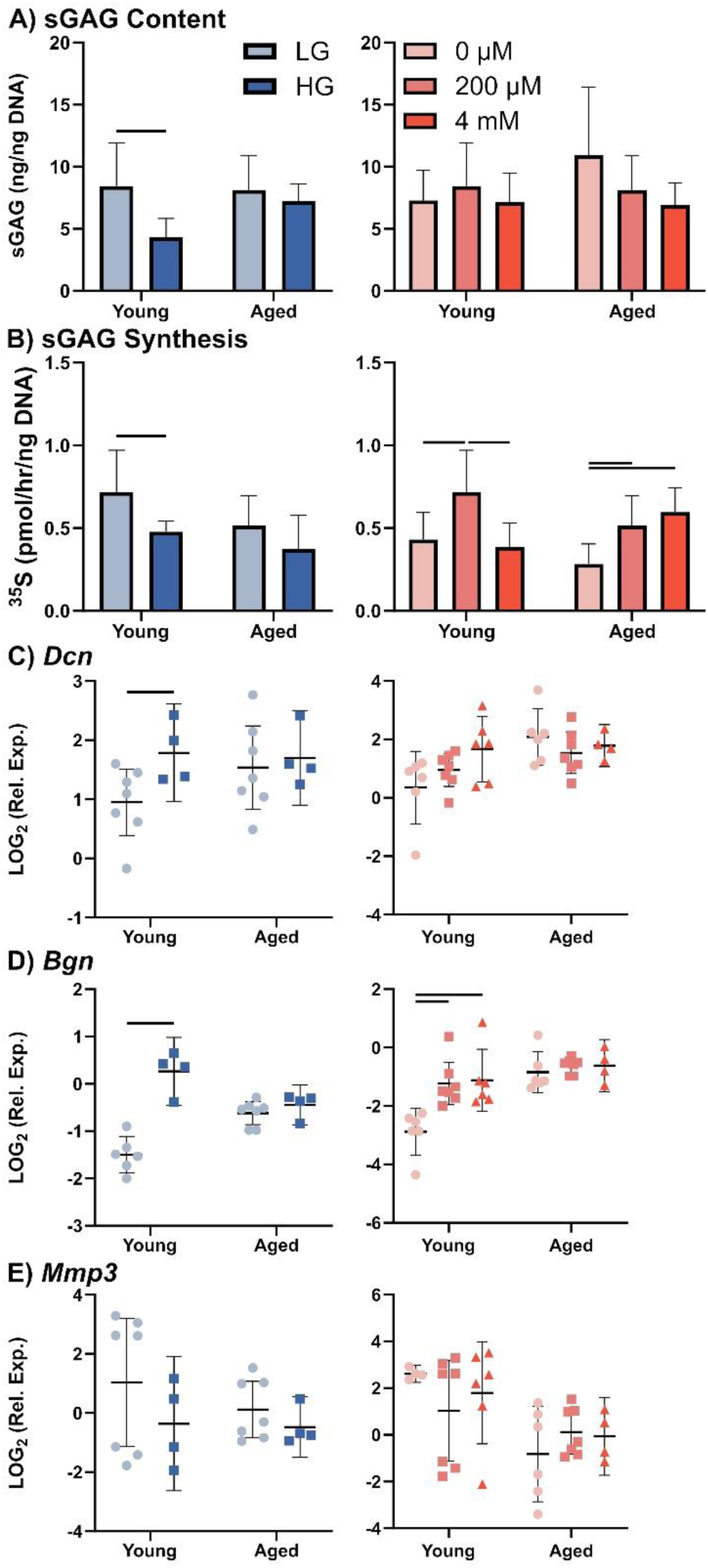
Measurement of (A) sGAG content, (B) sGAG synthesis, (C) decorin gene expression,(D) biglycan gene expression, and (E) MMP-3 gene expression for young and aged explants as a function of glucose concentration (left) and glutamine concentration (right). Data is presented as mean ± 95% confidence interval. Significance between media groups is denoted with a bar (-) with p<0.05. [LG = low glucose, HG = high glucose, 0 µM = 0 µM glutamine, 200 µM = 200 µM glutamine, 4 mM = 4 mM glutamine]

There were no significant changes in collagen content with either glucose or glutamine (Figure 5A). Collagen synthesis and collagen I gene expression, however, were stimulated by high glucose (Figure 5B and 5C). Counterintuitively, collagen synthesis was suppressed in the 4 mM glutamine group compared to the 0 and 200 µM groups in young tendons (Figure 5B).

**Figure 5.**
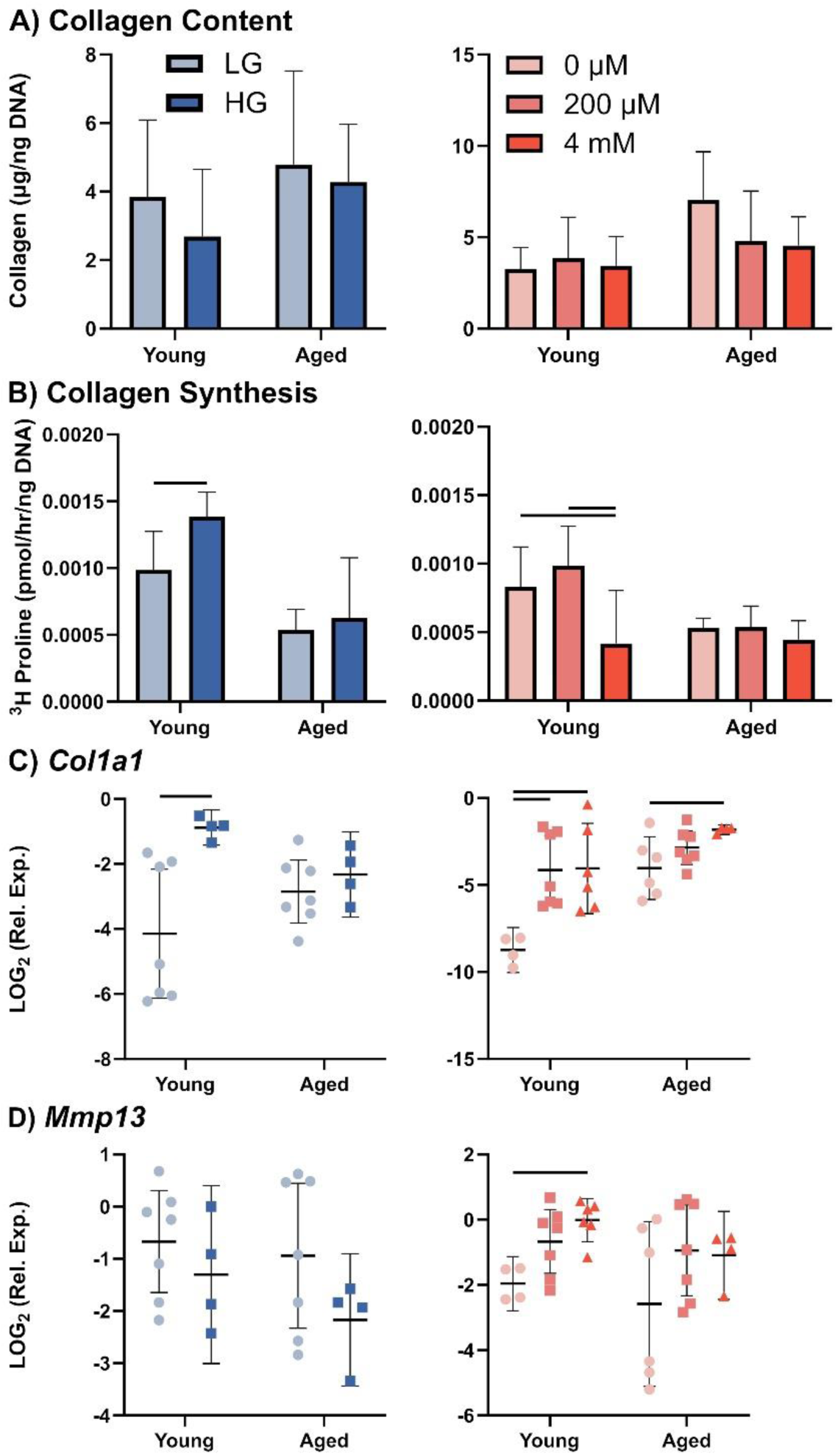
Measurement of (A) collagen content, (B) collagen synthesis, (C) collagen I gene expression, (D) MMP-13 gene expression for young and aged explants as a function of glucose concentration (left) and glutamine concentration (right). Data is presented as mean ± 95% confidence interval. Significance between media groups is denoted with a bar (-) with p<0.05. [LG = low glucose, HG = high glucose, 0 µM = 0 µM glutamine, 200 µM = 200 µM glutamine, 4 mM = 4 mM glutamine]

These changes were not replicated in aged groups. Collagen I gene expression, however, was upregulated by glutamine in both age groups (Figure 5C). Interestingly, the no glutamine group significantly reduced collagen I gene expression in young, but not in aged explants. Matrix metallopeptidase 13 (MMP-13) expression was also reduced in the no glutamine group, but only in young explants (Figure 5D).

### High glutamine causes inflammatory and apoptotic responses in aged tendons

Finally, we looked at how these changes affected markers of apoptosis and inflammation, since diabetes models and previous tenocyte studies had observed changes in similar markers with high glucose.^15,28^ Caspase-3 expression was upregulated by 4 mM glutamine in aged explants (Figure 6A). Gene expression of interleukin 6 (IL-6) was decreased by high glucose and increased by high glutamine in young explants (Figure 6B). And, in aged explants, high glutamine increased expression of tumor necrosis factor alpha (TNF-α) (Figure 6C).

**Figure 6.**
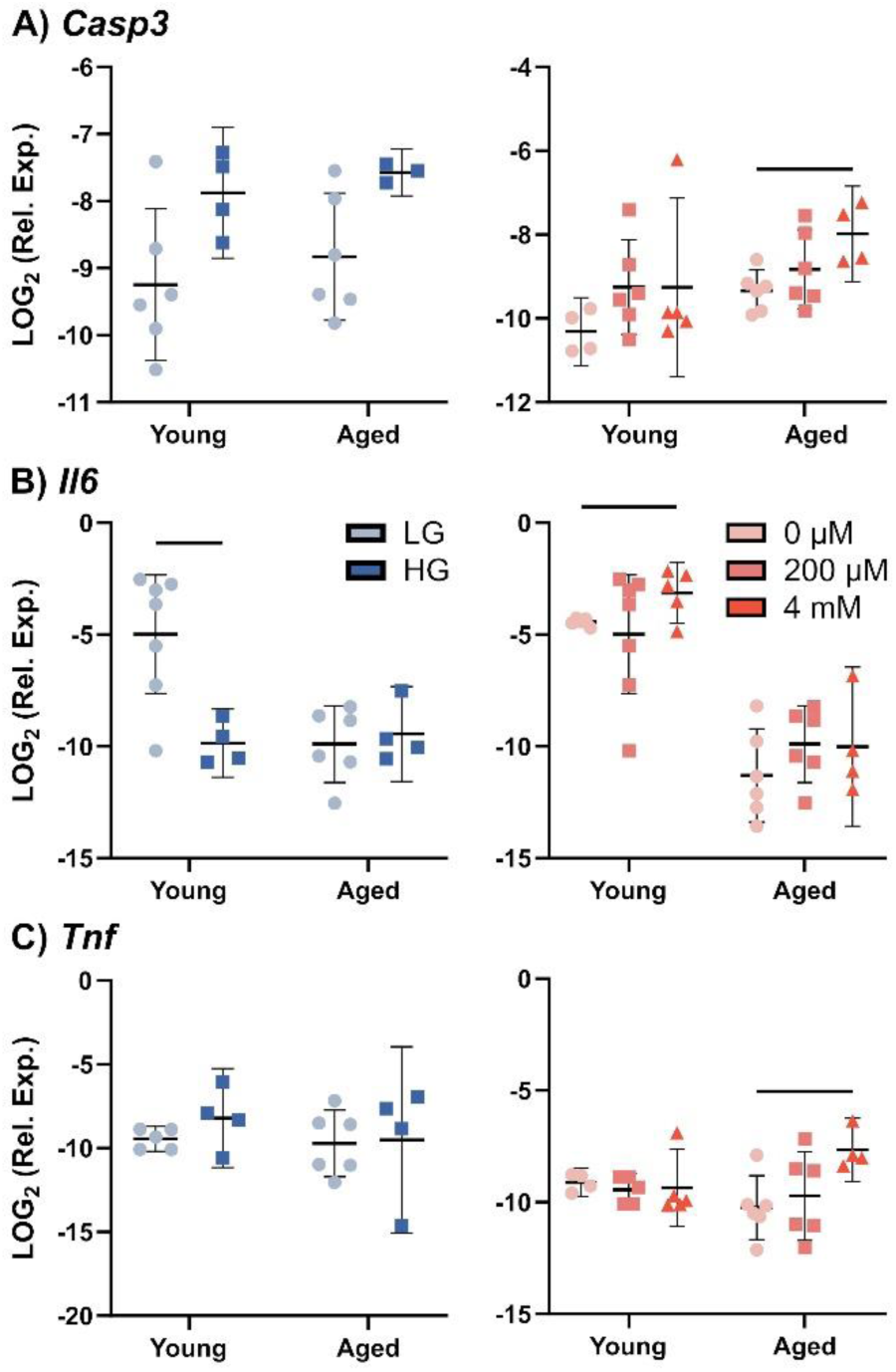
Measurement of mRNA levels of (A) CASP-3, (B) IL-6, and (C) TNF-α for young and aged explants as a function of glucose concentration (left) and glutamine concentration (right). Data is presented as mean ± 95% confidence interval. Significance between media conditions is denoted with a bar (-) with p<0.05. [LG = low glucose, HG = high glucose, 0 µM = 0 µM glutamine, 200 µM = 200 µM glutamine, 4 mM = 4 mM glutamine]

## Discussion

This is the first study to examine the coordination of glucose and glutamine supplementation in a tendon explant model. While some previous studies have looked at the individual effect of glucose or glutamine in tenocytes or *in vivo* models, these studies did not investigate the tissue-level response of tendon, the influence of aging, or the crosstalk between them. Our study interestingly found an age-dependent reliance of glutamine processing on glucose levels. Figure 7 shows the response of key enzymes and proteins in the glucose and glutamine pathways. Similar to previous studies, we found that glutamine uptake increased with high glucose, which could be the reason we see an increase in expression of most glutamine pathway genes with increased glucose concentration.^24,29^ Glutamine enters the cell through amino acid transporters, like SLC1A5, which we found to be upregulated with high glucose. The increase in SLC1A5 expression could be due to increased N-glycosylation due to increased glucose metabolism, which has been observed in previous studies.^30^ We also saw decreased SLC1A5 expression with increased glutamine. Since SLC1A5 is a glutamine transport protein, this could be a feedback mechanism for times when there is already enough glutamine inside the cell so less transport is required. Once glutamine enters the cell, it can be converted to glutamate and used for various biochemical functions, or it can enter the mitochondria, where it is mainly used as an energy source.

**Figure 7.**
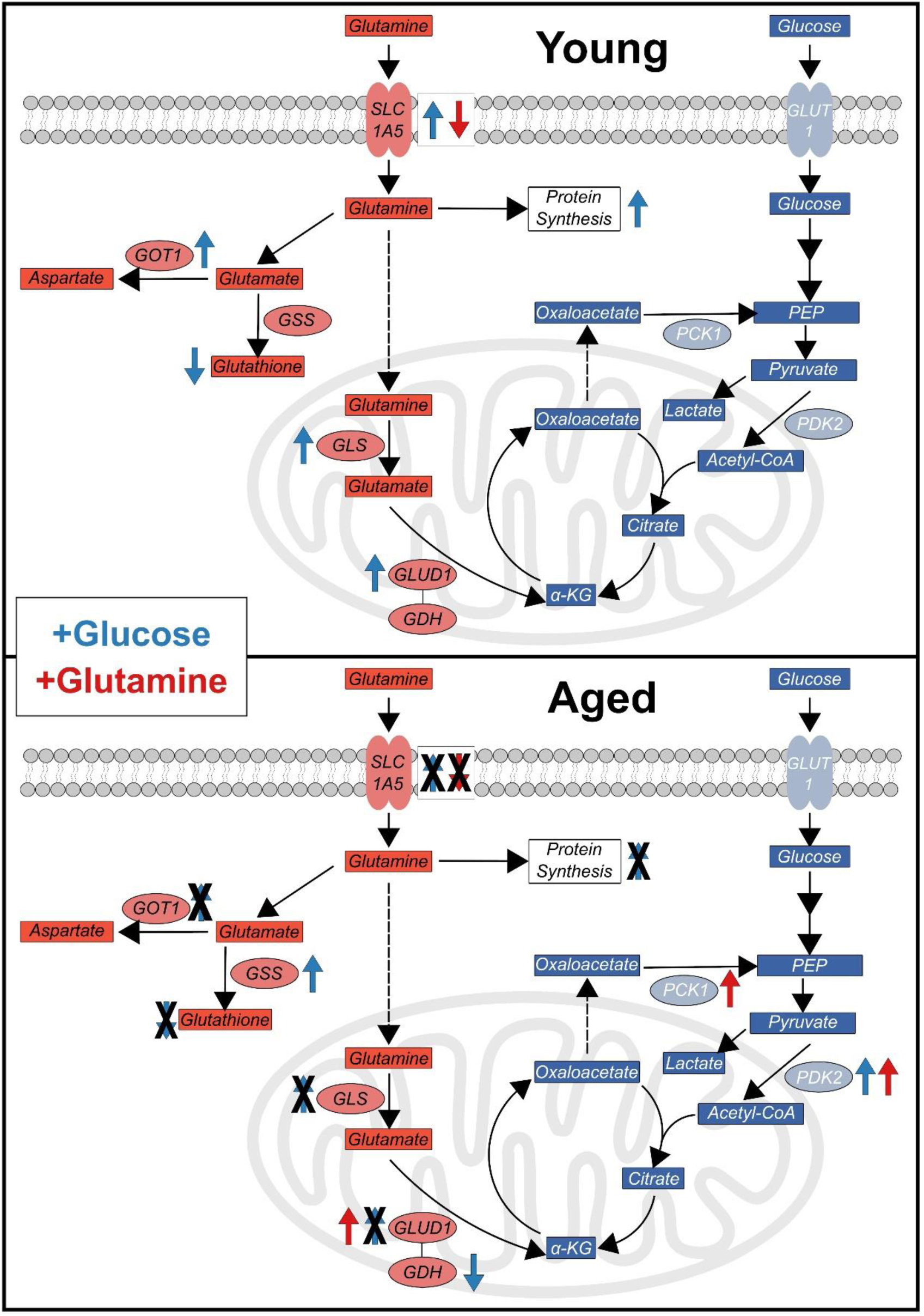
Summary of key points of the glucose and glutamine pathways and how their gene expression changes with increased glucose and glutamine content in young and aged explants.

Outside of the mitochondria, we found an upregulation of GOT1, an enzyme that can reversibly convert aspartate and α-ketoglutarate to glutamate and oxaloacetate. While the forward direction is more common, since this reaction can go in either direction, this result can be indicative of a few possibilities. In the case of the forward reaction, where glutamate and oxaloacetate are produced, this can be a sign of the activation of the malate-aspartate shuttle. In the malate-aspartate shuttle, oxaloacetate receives two electrons produced by glycolysis and moves across the mitochondrial membrane where it is converted back into oxaloacetate, so these electrons can be used for oxidative phosphorylation.^31^ Therefore, GOT1 could be upregulated due to the increased electrons produced by glycolysis in high glucose conditions. However, some studies have also observed the reverse reaction occurring, leading to the synthesis of aspartate, a non-essential amino acid used for a variety of purposes, including protein synthesis.^32,33^ We also see elevated protein synthesis in this study, supporting the possibility that there is significant aspartate made through GOT1 here, especially since in high glucose conditions cells may already have sufficient energy and don’t need the extra electrons from the forward reaction.

When glutamine instead enters the mitochondria, it is converted into glutamate by GLS, which we found to be upregulated in young tendons with high glucose. This glutamate can then be converted into α-ketoglutarate by GDH, which is encoded by GLUD1, and can then be used in the citric acid (TCA) cycle. Since these proteins are used to create TCA cycle intermediates, it makes sense that they would be upregulated in response to high glucose. In aged animals, we instead see a decrease in GDH content, signifying a shift in glutamine catabolism with aging.

This change is likely due to age-related changes in glucose metabolism, since GDH is inhibited by ATP, which decreases with age.^34^ Since GDH requires NAD+ to convert glutamate, these changes could also be due to decreased NAD+ levels that have previously been observed in aging or due to high glucose.^35,36^ Similar to other studies, we also see an increase in baseline GDH activity with age.^37,38^ Aging could affect the body’s metabolic demands, so more glutamine is converted to α-ketoglutarate for use in TCA, but higher glucose in aged reverses this effect since it is unnecessary.

We also observed an age-dependent response in glutathione (GSH), which helps protects cells from reactive oxygen species (ROS). We found that GSH decreased with high glucose in young, which has previously been observed in epithelial and vascular smooth muscle cells.^39,40^ This change could be due to high glucose-induced oxidative stress depleting GSH.^41,42^ We don’t see the same response in aged explants, but it is possible that our assay isn’t sensitive enough to pick up this change due to the lower baseline concentration in aged explants. The baseline lower GSH content in aged samples has been observed in many other tissues and is an example of age-related changes in glutamate catabolism.^43^ Previous studies have observed similar changes in glutamine metabolism with age due to aging-related mitochondrial dysfunction where the catabolism of glutamate is altered, shifting towards different pathways compared to healthy mitochondria.^44^ There also seems to be a tradeoff between GSH and GDH with age, with more conversion of glutamine to GSH in young samples and more to GDH in aged samples. Future studies should investigate conversion of glutamine to other end products, like alanine and aspartate, to examine how catabolism changes with aging.

We also observed age-dependent changes in glucose processing markers. Contrary to the glutamine pathways, however, we only saw changes in glucose processing in aged tendons. PDK2 expression increased with both high glucose and our highest glutamine concentration. PDK2 inhibits pyruvate dehydrogenase, which converts pyruvate into acetyl-CoA, so an increase in PDK2 could signal a shift from energy production via oxidative phosphorylation in the TCA cycle to glycolysis with pyruvate being converted into lactate instead.^45^ This shift with aging could be a sign of metabolic stress or impaired glucose tolerance.^46,47^ PDK2 is also inhibited by ROS, which are increased in aged tissue.^48^ Therefore, this could be an adaptive response to limit the influx of pyruvate and decrease production of ROS. In addition, we saw an increase in PCK1 expression in our highest glutamine concentration in aged explants. PCK1 is responsible for converting oxaloacetate, from the TCA cycle, back into phosphoenolpyruvate, which can then be converted back into pyruvate to generate ATP in glycolysis. Aging is associated with a shift to using glutamine to create TCA cycle intermediates due to increased energy needs from mitochondrial dysfunction, so this increase in expression could be in response to increased oxaloacetate from the TCA cycle.^44^ Overexpression of PDK2 and PCK1 also play a key role in both cancer and diabetes, so these increases in aged tendons are likely maladaptive.^49–52^

In general, Figure 7 shows a shift from high glucose regulation of the glutamine pathway in young tendons to regulation of the glucose pathway by high glutamine in aged tendons. While many possible explanations have been proposed, the exact mechanism by which glucose regulates the glutamine pathway, and vice versa, is unknown.^24^ The glutamine pathway is vital in cell and tissue homeostasis, serving a vital role in protein synthesis, redox homeostasis via ROS removal, and nucleotide synthesis. When there is excess glucose in our young tendons, we see an increase in protein synthesis, signifying remodeling to allow tendon to maintain homeostasis.

However, this does not happen in our aged tendons, which could signify a breakdown of the signaling mechanism between glucose and glutamine or increased need of glucose for energy production rather than for use in signaling with aging. This lack of signaling could potentially be one of the main causes for aged tendons’ reduced mechanical properties and impaired regenerative potential post-injury.^53,54^

Furthermore, our study also demonstrates that these key metabolites influence ECM homeostasis. In young explants we saw an increase in collagen synthesis and collagen I gene expression with high glucose. We also found increases in metabolism which could lead to more synthesis, especially given that synthesis is normalized to cell count and thus represents synthesis per cell. However, this increased collagen synthesis and our observed increase in collagen I gene expression with high glucose do not result in increased collagen content, suggesting degradative activity that might balance increased synthesis. We did not find an increase in MMP-13 gene levels, so this degradation is likely due to some other MMP or cathepsin. Our results disagree with previous studies in tenocytes and tendon-derived stem cells which found decreases in collagen I expression due to high glucose.^14,17^ These studies’ measurements were performed at 48 hours, while ours was at 7 days, so there could be a difference in the timing of the two responses. This difference could also be a result of our explant model system, which is more physiologically relevant than isolated 2D cell studies. Furthermore, these findings could be dependent on other environmental factors, such as oxygen level or temperature.^55^ Future studies will focus on earlier and later timepoints to see how this response is altered over time. Interestingly, our highest glutamine level also led to decreased collagen synthesis. Previous studies found the opposite response with increased collagen synthesis with increased glutamine in other tissues.^18,22^ It is possible that this glutamine level is too high and potentially leads to some kind of maladaptive response.

We also see age-specific changes in other key ECM proteins, proteoglycans, because of medium glucose and glutamine concentrations. First, we found that high glucose suppressed sGAG synthesis and content in young tendons, consistent with other studies in tendon and other tissues.^56,57^ This decrease in content and synthesis was surprisingly accompanied by an increase in decorin and biglycan gene expression. Our results disagree with those of previous tenocyte studies, which found no change in biglycan and decorin at day 7, possibly due to our study controlling for glutamine concentration or due to the influence of using an intact tendon in our explant model.^16^ Glutamine concentration also affected sGAG synthesis with 200 μM being best in young and 4 mM being best in aged, meaning it is likely that glutamine leads to an increase in amino acids required for sGAG formation. Since sGAG synthesis is reliant on both glucose and glutamine levels, it hints at a possible molecular link between the two pathways. This link could potentially be found in the hexosamine biosynthesis pathway, which is a key component of proteoglycans, and requires input from both glucose and glutamine pathways.^24,29^

Finally, we sought to investigate how glucose and glutamine concentrations mediate changes in inflammation markers, since hyperglycemia and diabetes generally cause an increase in inflammation.^58,59^ We found that IL-6 decreases with high glucose levels in young explants, but since this is not accompanied by increased TNF-α, it is likely that this is a signaling and not an inflammatory response. IL-6 regulates glucose metabolism with increased levels of IL-6 being associated with increased ability to metabolize glucose.^60^ It’s possibly downregulated here to limit glucose uptake in an environment with excess glucose. Similar results were found in muscle where endurance exercise-induced IL-6 is downregulated by glucose ingestion.^61^ We also saw an upregulation of TNF-α with our highest level of glutamine in aged tendons. Previous research found that chronically painful tendons have a higher concentration of glutamate than pain-free tendon, which means that this results could be a sign of a stress response caused by excessive glutamine.^62,63^ This stress response caused by our highest level of glutamine in aged explants is further supported by an upregulation of caspase-3, indicative of elevated apoptosis.

This response could be caused by an increase in ROS or ammonia, which aged explants cannot process as effectively due to lower GSH production.^37,64^ While negative effects from excess glutamine have been observed in several major organs and bone, this is the first study to observe a similar stress response in tendon.^65–67^ This maladaptive response to a 4 mM concentration of glutamine, the most common concentration in most media, supports that *in vitro* studies should be more careful with controlling glutamine concentration to maintain tendon health.

Our model has a few limitations that will need to be further investigated. First, we culture our tendons under stress-deprivation rather than with a tensile static or cyclic load that would better mimic physiologic conditions. Our previous studies have demonstrated no changes in tendon health in stress deprivation over the culture period studied here. However, future studies will look to decouple effects related to mechanical loading and changes to glucose and glutamine metabolism through loading experiments.^2^ Also, this study measured changes at a single timepoint. We plan to investigate both earlier and later timepoints to identify mechanisms of dysregulation and potential targets for reversing maladaptation. Our study used a targeted approach to reveal changes in specific genes and proteins within the metabolic pathways, but future studies will capitalize on more broad-spectrum approaches such as metabolomics and lipidomics. Finally, we didn’t investigate any of the potential pathways that cause crosstalk between glucose and glutamine or quantify reactive oxygen species; future studies will investigate signaling pathways, like mTOR and AMPK, to further understand these interactions.

Overall, this work reveals age-dependent changes in tendon explants to various concentrations of glucose and glutamine which likely have a significant influence on ECM remodeling and tissue health. Importantly, we found that glutamine and glucose processing is disrupted with aging. Given the importance of glucose and glutamine metabolism on the synthesis and degradation of extracellular matrix proteins, we believe it is vital to understand how age-related changes to these processes affect tissue homeostasis and disease progression in other tissues as well. This study suggests that age-related metabolic changes, like mitochondrial dysfunction, impaired glucose tolerance, and metabolic stress, could be some of the main factors that promote tendon degeneration in aged populations. If we can identify the specific changes, we will be able to find targets to prevent or reverse age-related dysfunction and thus halt degeneration early.

## Acknowledgements

This study was supported by Boston University and NIH/NIA R00-AG063896. We would also like to thank Emily Larson and Brandon Kao for contributions in the early development of this project.

## Author contributions

S.J.M. and F.R.P. have contributed to all aspects of this study, including research design, data acquisition, interpretation/analysis of data, and drafting/revision of the manuscript. K.J.S. has contributed significantly to research design, interpretation/analysis of data, and drafting/revision of the manuscript. B.K.C. has contributed significantly to research design, interpretation/analysis of data, and drafting/revision of the manuscript. All authors have read and approved the final submitted manuscript.

## Methods

### Study Groups and Tissue Harvest

Flexor digitorum longus (FDL) tendon explants were harvested from the limbs of young (4 months) and aged (20 months) C57BL/6J male mice from JAX. Our young group represents a human age equivalent of 20-30 years, which is considered skeletally mature, and our aged group represent a human age equivalent of 55-65 years of age, a clinically relevant population for tendon degeneration^68^. Tendons were harvested directly following sacrifice using previously described methods^69^ per approved animal use protocol (BU IACUC PROTO202000046). The proximal and distal ends of the tendons were trimmed to only include a 10-mm intrasynovial segment of the FDL for explant culture.

### Explant Culture

Explants were cultured in stress-deprived conditions (no mechanical stimulus) submerged in modified culture medium. Low glucose (1 g/L; Lonza) and high glucose (4.5 g/L; Cytiva) Dulbecco’s Modified Eagle’s Media without glutamine was supplemented with 10% fetal bovine serum (Cytiva), 100 units/mL penicillin G (Fisher Scientific), 100 μg/mL streptomycin (Fisher Scientific), 0.25 µg/mL Amphotericin B (Sigma-Aldrich). Variable amounts of L-glutamine solution (200 mM; Fisher Scientific) were added for final concentrations of 0 µM, 200 µM, and 4 mM of glutamine in medium. Medium was made fresh to limit the degradation of L-glutamine^70^ and changed every 2 days. After 7 days, explants were collected to assess changes in biosynthesis, composition, metabolism, and gene expression. Spent medium was also collected periodically to measure uptake of glucose and glutamine.

### Metabolite Analyses

Levels of glucose and glutamine were determined via analysis of spent culture medium (n=4/group) using a Glucose Assay Kit and a Glutamine Assay Kit (Abcam), respectively. Medium glucose and glutamine levels were subtracted from recorded levels in wells with just cultured medium with no tendons to measure change in metabolite concentration. A positive value represented a decrease of metabolite in the medium, meaning the tissue was taking up and processing the metabolite. A value of zero represented no difference in metabolite level between medium and medium containing tissue, suggesting no influx or efflux. A negative value represented an increase of metabolite in the medium, suggesting production of the metabolite by the tissue. To measure protein content of key glutamine enzymes, tendons were first digested and then levels of glutathione protein content were measured using a GSH-Glo assay (Promega) and glutamate dehydrogenase content was measured using a glutamate dehydrogenase activity assay kit (Abcam) following manufacturer’s protocols.

### Gene Expression

Explants were collected from each medium group at day 7, flash frozen in liquid nitrogen, and stored at -80°C for analysis of gene expression (n=3-7/group). Samples were homogenized in TRIzol reagent with a bead homogenizer (Benchmark Scientific), and then separated using phase-gel tubes (Qiagen). The supernatant was then purified following the Quick-RNA MicroPrep Kit protocol (Zymo Research) and the quantity and quality of RNA was verified on a microplate reader (Agilent). Total RNA was reverse transcribed, and RT-PCR was performed using the Applied Biosystems StepOne Plus RT-PCR (Applied Biosystems) with SYBR Green Master Mix (Fisher Scientific). Primer pairs and sequences are listed in the Supplemental Data (Table S1). Genes targeted were involved in glucose metabolism (*Glut1*, *Pdk2*, *Pck1*), glutamine metabolism (*Slc1a5*, *Glud1*, *Got1, Gls, Gss*), matrix synthesis (*Col1a1*), regulation of collagen fibrillogenesis *(Dcn*, *Bgn*), matrix degradation (*Mmp3*, *Mmp13*), and injury (*Il6*, *Tnfa, Casp3*). Expression for each gene was calculated from the threshold cycle (Ct) value and was normalized to the housekeeping gene β-Actin. All data are represented in log space.

### Biosynthesis and Composition

Explant cell metabolism (mitochondrial respiration) was measured using the resazurin reduction assay, as previously described.^71^ Following a 3-hour incubation of resazurin solution in culture medium, intensity of the reduced product, resorufin, was measured using excitation/emission of 554/586 nm on a microplate reader. Values were normalized to daily control wells with no explant, such that a value below 1 was representative of a nonviable explant. Following culture, explants (n = 7-12/group) were washed for 2 hours and then digested with proteinase K (5 mg/mL) (Sigma-Aldrich) for 16 hours. Synthesis of sulfated glycosaminoglycans (sGAG) and total protein (indicative of collagen synthesis) was measured by 24-hour incorporation of ^35^S-sulfate (20 μCi/mL) and ^3^H-proline (5 μCi/mL), respectively (Perkin-Elmer). Radiolabel incorporation in digested tissue was measured with liquid scintillation counter (Perkin-Elmer).^4^ Double-stranded DNA content was measured using the PicoGreen dye binding assay.^72^ sGAG content of the tissue digest was measured using the dimethyl methylene blue (DMMB) assay.^73^ Then 100μL of each tissue digest was hydrolyzed using 12 M HCl, dried, resuspended, and assayed to measure total collagen content using the hydroxyproline (OHP) assay.^74^ Biochemical data (total protein synthesis, sGAG synthesis, sGAG content, collagen content) are presented normalized to DNA content.

### Statistics

Data points more than two standard deviations outside of the mean were removed as outliers, which resulted in two samples in separate groups being removed from our PCR data. Statistical evaluations for glucose and glutamine uptake were performed using two-way ANOVA (effects of group and age). Statistical evaluations comparing across glutamine concentration were performed with a one-way ANOVAs while evaluations comparing across glucose concentrations were performed with t-tests within each age group. All ANOVAs were performed with Bonferroni corrected t-tests and significance was p < 0.05 for all tests. All data are presented as mean ± 95% confidence interval.

