## Supplemental Table S1 for "Coordination of Glucose and Glutamine Metabolism in Tendon is Lost in Aging"

**Supplemental Table S1.** PCR primer forward and reverse sequences used for quantitative gene expression analysis.

| **Gene Name** | **Forward (5’→ 3’)** | **Reverse (5’→ 3’)** |
| --- | --- | --- |
| *Actb*^1^ | GGCTGTATTCCCCTCCATCG | CCAGTTGGTAACAATGCCATGT |
| *Glut1* | CCCAGAAGGTAAGTGTGCTATG | GCTCCTCTTCTCTGACCATTTC |
| *Pdk2* | GACTGCCAAGACAGGCTAAA | GCTGGTGTGAAGGAGAGTAATG |
| *Pck1* | CCGAGACAAGTCTATGGGTAAAG | GCCAGAGAAGGAGGTATGAATG |
| *Gls*^2^ | CATCCTCATCTGACGAGCGG | TCCTGTAGGATCTCCGAGGG |
| *Gss* | CCATGGGCTACTCTAACCATTC | GCACGATGACTCACACCTATAC |
| *Glud1* | CATCAACTCGCTACTCCATTCT | CCACCCATATCTGACCCTAAAC |
| *Slc1a5* | AGAGATGAGGCTCCTGGATAA | GTGGGAAGGGTACAAGGATAAA |
| *Got1* | CTAGCCGCCTCCCTTTATTT | CATTGCCGGACGATCCTATTA |
| *Dcn*^3^ | CTATGTGCCCCTACCGATGC | CAGAACACTGCACCACTCGAAG |
| *Bgn* | TTTCTGAGCTTCGCAAGGATG | GGGCGTAGAGGTGCTGGAG |
| *Mmp3*^3^ | ACATGGAGACTTTGTCCCTTTTG | TTGGCTGAGTGGTAGAGTCCC |
| *Col1a1*^3^ | GACATGTTCAGCTTTGTGGACCTC | GGGACCCTTAGGCCATTGTGTA |
| *Mmp13*^4^ | TCAGTCTCTTCACCTCTTTTGGGAATCC | TCAGTTTCTTTATGGTCCAGGCGATG |

1.	Veres-Székely A, Pap D, Sziksz E, et al. 2017. Selective measurement of α smooth muscle actin: why β-actin can not be used as a housekeeping gene when tissue fibrosis occurs. BMC Mol. Biol. 18(1):12.

2.	Ke C, Gao J, Tu J, et al. 2022. Ganfule capsule alleviates bile duct ligation-induced liver fibrosis in mice by inhibiting glutamine metabolism. Front. Pharmacol. 13:930785.

3.	Connizzo BK, Piet JM, Shefelbine SJ, Grodzinsky AJ. 2020. Age-associated changes in the response of tendon explants to stress deprivation is sex-dependent. Connect. Tissue Res. 61(1):48–62.

4.	Connizzo BK, Grodzinsky AJ. 2018. Release of pro-inflammatory cytokines from muscle and bone causes tenocyte death in a novel rotator cuff in vitro explant culture model. Connect. Tissue Res. 59(5):423–436.
