## Supplementary figures and images for "Coordination of Glucose and Glutamine Metabolism in Tendon is Lost in Aging"

### Supplemental Figure S1

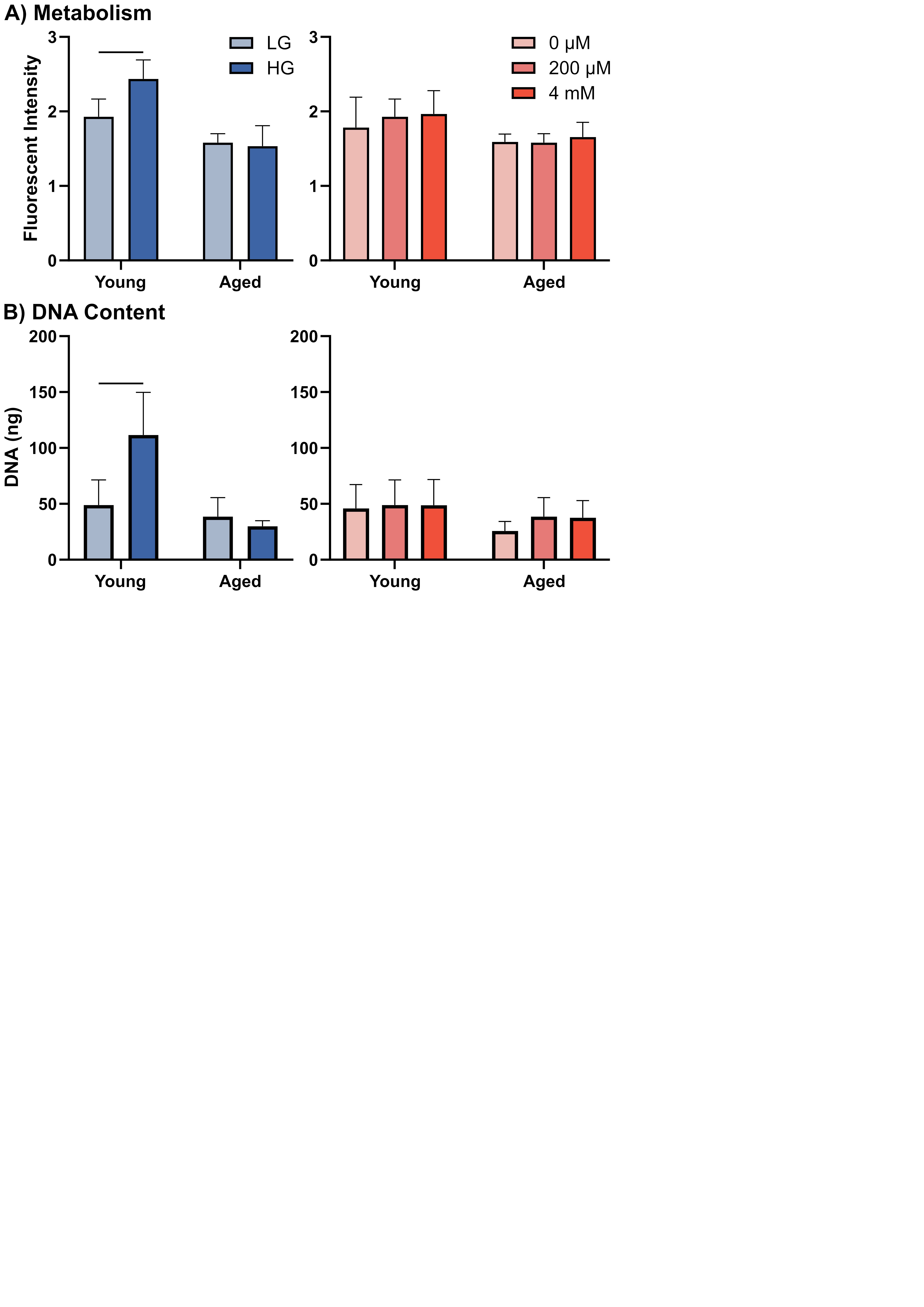

### Supplemental Figure S2

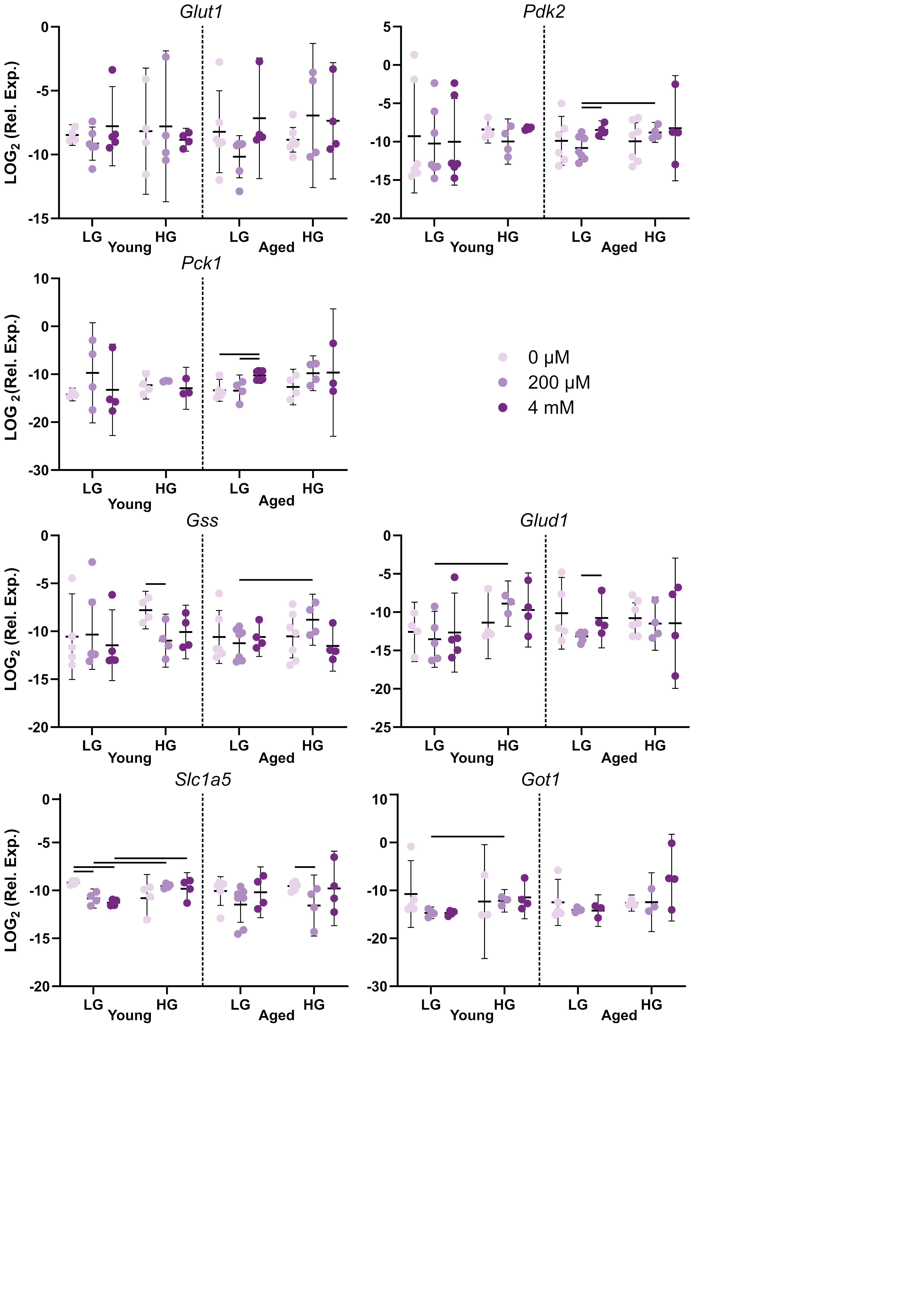

### Supplemental Figure S3

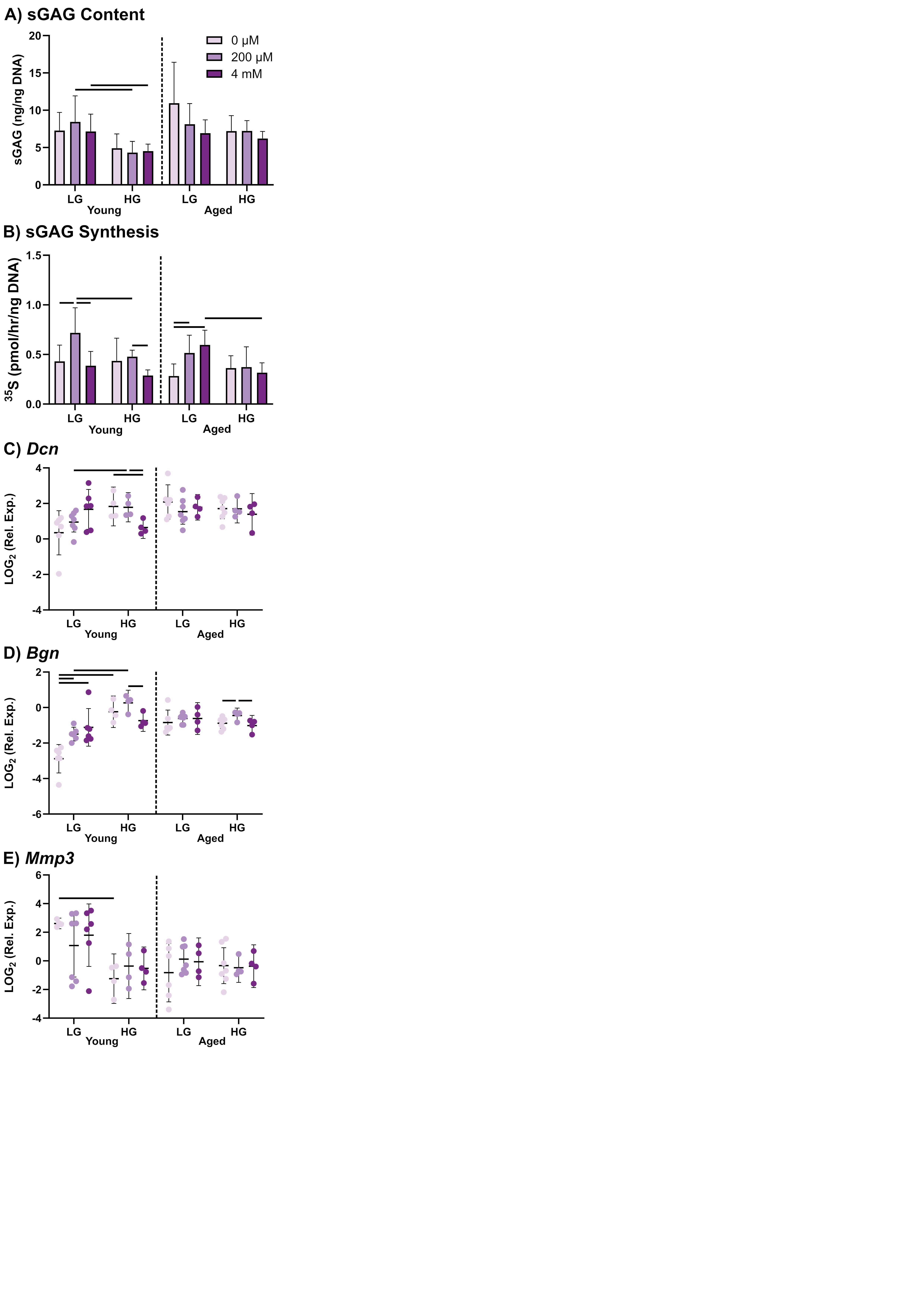

### Supplemental Figure S4

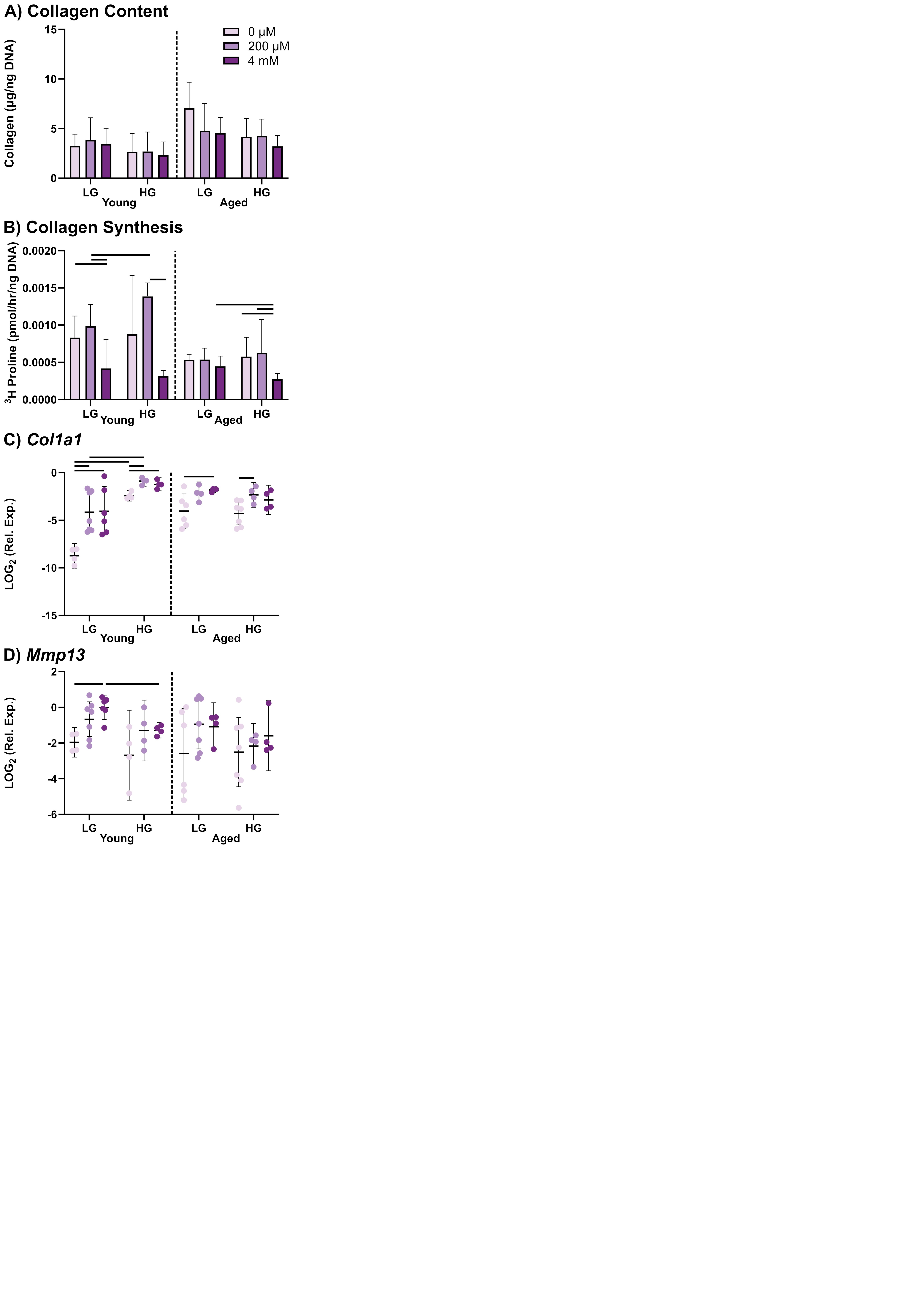

### Supplemental Figure S5

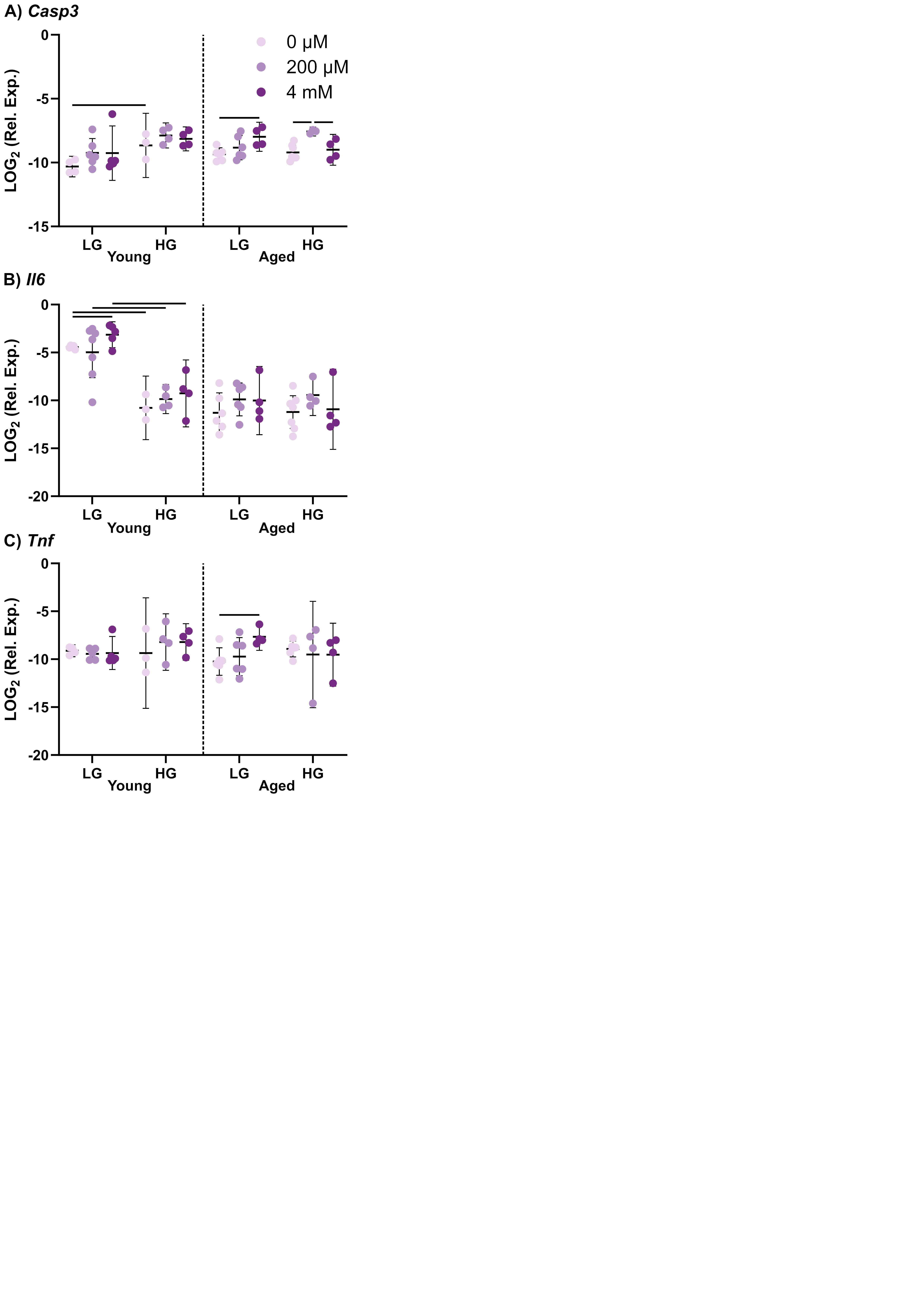
